# From Shadows to Data: A Robust Population Assessment of Snow Leopards in the Highland Crossroads

**DOI:** 10.1101/2025.03.26.645478

**Authors:** Muhammad Ali Nawaz, Shoaib Hameed, Jaffar Ud Din, Hussain Ali, Shakeel Ahmad, Ian Durbatch, Mehmood Ghaznavi, Mohsin Farooque, Naeem Iftikhar, Muhammad Samar Hussain Khan

**Author notes:** **Corresponding Author**: Muhammad Ali Nawaz, **Email id**.

## Abstract

The snow leopard (*Panthera uncia*) is a flagship species of the greater Himalayan region - referred to as the ‘Third Pole’ - and symbolizes integrity of this ecological system. Within the greater Himalayas, Pakistan holds special significance as the north of the country represents a confluence of four major mountain ranges (Hindu Kush, Pamir, Karakoram, and Himalaya). However, robustly surveying and monitoring elusive, low-density species such as snow leopards has historically been difficult in the region. As a result, our understanding of the spatial patterns in density and overall population size of snow leopards has remained conjectural in the ‘highland crossroads’ of northern Pakistan. This lack of objective information is an obstacle to realizing effective conservation planning for the species in Pakistan, as well as the broader ecosystem within which it plays a key role. This study aimed to empirically derive population estimates for snow leopards in Pakistan, based on robust camera trapping. Extensive camera trapping was conducted covering about 39% of the snow leopard range in Pakistan from 2010 to 2019, spread across the four major mountain ranges in the north of the country. A total of 828 cameras were placed over 26,540 trap days, resulting in 4,712 photos of snow leopards obtained from 65 different locations. Among the 53 unique individuals identified, the majority (53%) were detected only once, with an overall recapture frequency of 2.28 times per individual. Spatial capture-recapture (SCR) was employed for population and density estimation. Model selection strongly favored a model in which density was positively associated with elevation, and camera type influenced baseline encounter rates. The estimated population size for snow leopards in this highland crossroads was 127 (95% CI 88-182) adult animals, with a mean density of 0.13 (95% CI 0.09-0.19) animals per 100 km². Examination of the density predictions revealed that higher density areas were associated with protected areas and greater prey biomass, highlighting the importance of these two key factors. This research provides the first robust population estimate for snow leopards in this region, establishing a foundation for long-term population monitoring and assessing the effectiveness of conservation measures. We recommend the integration of complementary approaches, such as non-invasive genetic methods, to validate and refine population estimates.

## Introduction

The greater Himalayan region, referred to as the ‘Third Pole’ due to its towering mountains and massive glacier deposits, is a critical global ecosystem. Rich in natural resources, it supports about 20% of the world’s human population in terms of water and associated ecosystem services and serves as a global biodiversity hotspot. Northern Pakistan holds special significance within this region as it is the confluence of four major mountain ranges that define this region: the Hindu Kush, Pamir, Karakoram, and Himalayas. This makes the area ecologically unique, offering a diverse assemblage of species and serving as a crucial connectivity corridor for wide-ranging species.

To underpin effective conservation strategies for the region, accurate species monitoring is required. However, species monitoring, especially for threatened species, remains an important challenge in modern conservation biology and resource management (Gopalaswamy et al. 2019, Elliot et al. 2020). This challenge is particularly acute for large carnivores, which typically have large home ranges, exist at low densities, and are often nocturnal and camouflaged (Crooks et al. 2011, Trouwborst 2015). Additionally, some large carnivore species inhabit densely vegetated areas or occupy remote and inaccessible high-altitude regions, complicating density estimation financially, logistically, and methodologically (Jackson et al. 2006, Watts et al. 2019, Janjua et al. 2020).

Technological advancements in noninvasive survey techniques, such as molecular genetics and camera traps, along with statistical tools to model imperfect detection—such as spatial capture-recapture (SCR) (Borchers and Efford 2008, Royle et al. 2014)—have revolutionized wildlife monitoring. These methods provide the means to acquire data at large scales and the framework to generate spatially explicit predictions (Linden et al. 2020). Individual-based identification is crucial for efficient and reliable population estimation of large carnivores that comprise a wild population (Balme et al. 2019). Identification from remote cameras, particularly for species with unique coat marking such as snow leopards (*Panthera uncia*), enables sampling across extensive areas. Detections on cameras allows for the creation of individual capture histories to estimate population size and density at both local and global scales ( Royle et al., 2013; Johansson et al., 2020).

The snow leopard, a solitary apex predator, inhabits high-altitude mountainous regions across South and Central Asia. Its distribution spans highly rugged mountain ranges such as the Altai, Pamir, Tien Shan, Hindu Kush, Himalayas, and Karakoram (McCarthy and Chapron 2014, McCarthy et al. 2017). The species serves as an indicator of the health and biological integrity of this high mountain ecosystem (Sharief et al. 2022).

Despite its ecological significance, the snow leopard faces multiple threats throughout its range, including prey depletion, poaching, illegal trade, retaliatory killings due to livestock predation, climate change, and habitat loss due to infrastructure development (McCarthy et al. 2017, Ale and Mishra 2018, Li et al. 2020, Shen 2020, Din et al. 2022a). Consequently, the species is classified as Vulnerable on the IUCN Red List (IUCN, 2024) and placed under Appendix I of CITES, which prohibits the legal trade of its body parts and derivatives. It is also listed in Appendix I of the Convention on the Conservation of Migratory Species of Wild Animals (CMS), which requires immediate protection for the species by the parties.

Although snow leopards occupy a vast range (estimated at 2.8 million km² across 12 countries), their global population is relatively small, estimated at 3,900–8,745 individuals (GSLEP, 2024). These figures are largely derived from subjective assessments, as less than 3% of the global range has been surveyed using reliable techniques such as camera trapping or molecular genetics (Janečka et al. 2008, Chetri et al. 2019). The available data are further skewed toward high-density areas, creating a biased understanding of broader population sizes (Suryawanshi et al. 2019).

The paucity of robust data often necessitates reliance on expert opinion, which risks introducing inaccuracies and potentially harmful management decisions (Weise et al. 2017, Darimont et al. 2018, Moqanaki et al. 2018). Most of the current snow leopard population estimates rely on indirect methods, including interviews, questionnaire surveys, and sign-based evidence (e.g., tracks or scat), and thus lack the precision necessary for scientifically robust population estimates (Jones et al. 2008, Janečka et al. 2011).

In 2017, based on such subjective data, the IUCN downlisted the snow leopard’s status from Endangered to Vulnerable, assuming populations were larger than previously thought (McCarthy et al., 2017). This decision sparked controversy among scientists and governments, as it lacked support from empirical data (Ale and Mishra, 2018; GSLEP Secretariat, 2017). The consensus of the conservation community is essential for collective action, which can be achieved by utilizing data collected through scientifically robust methods that are agreed upon and owned by the stakeholders. Recognizing this need, the Global Snow Leopard and Ecosystem Protection Program (GSLEP) launched the Population Assessment of the World’s Snow Leopards (PAWS) initiative in 2017. This ambitious effort has facilitated the adoption of cutting-edge techniques for abundance estimation, enabling many snow leopard range countries to produce their first robust and replicable national population estimates for snow leopards.

The snow leopard is a national icon in Pakistan, with a distribution range of 80,000 km² across three major mountain ranges: the Hindu Kush, Himalayas, and Pamir-Karakoram (Hameed et al. 2020). Nevertheless, like other range countries, Pakistan observes a significant knowledge gap in species’ population ecology (Din et al. 2023). Historical assessments of the snow leopard population in Pakistan are largely speculative, relying on indirect methods and expert judgments. For instance, early estimates indicated a population of 100 (Anon, 1972), and 250 individuals (Schaller, 1976), based on surveys of 3,000 km². Malik (1997) suggested a population of 400 without specifying the surveyed area and Hussain (2003) reported 300–400 individuals based on questionnaire surveys covering 3,150 km². These “guesstimates” are extrapolated from less than 5% of the species’ range in Pakistan and lack true detections of the species. This limitation is understandable, as technology to aid in the monitoring of elusive large carnivores like snow leopards was at a far less advanced state than today.

To address this critical knowledge gap, our team, in collaboration with provincial/territorial wildlife departments, initiated extensive camera trapping studies in 2010. Given the challenging terrain, vast range, and logistical constraints, this effort required sustained dedication to achieve adequate coverage of the snow leopard’s presumed habitat. Over a period of 10 years, our surveys encompassed approximately 39% of Pakistan’s snow leopard range, yielding an unprecedented dataset of verifiable and replicable detections of the species.

This paper presents the first empirically derived population estimates for snow leopards in Pakistan, based on robust camera-trapping methodologies. The findings not only fill a crucial gap in snow leopard biology but also provide essential data to inform conservation planning in the region. Furthermore, this study offers valuable insights into the pitfalls and best practices of camera trapping in the high-altitude, rugged terrain of northern Pakistan, based on lessons learned from extensive camera trapping. This is expected to improve monitoring practices for large carnivores across High Asia and enhance the effectiveness of conservation efforts.

## Materials and Methods

### Study Area

The present study was conducted across the geographic range of snow leopards in Pakistan, spanning four major mountainous ranges: Hindu Kush, Himalaya, and Pamir-Karakoram (**Figure 1**). The area connects with China to the north, Afghanistan to the west, and India to the east. The elevation ranges from 566 to 8,570 m. Topography is characterized by high-altitude, snow-covered mountains, steep and high peaks, extremely rugged terrain, and highland plateaus. Climatic conditions vary widely across the study area, ranging from cold, semi-arid deserts to a monsoon-influenced, moist temperate zone (Sayed et al. 2013, Ahmad et al. 2022). The climate in the study area varies with altitude: winters are dry and cold, with temperatures falling below −20°C in the higher altitude areas (Zaman et al. 2024); summers are mild with temperatures rising to approximately 27 °C in May (Ali et al. 2017). Precipitation is mostly (90%) in the form of snow and ranges from 200 to 900 mm annually (Shedayi et al. 2016). Four vegetation zones are identified along altitudinal gradients: alpine meadows, sub-alpine scrub zones, alpine dry steppes, and permanent snowfields (Hameed et al. 2020). Mammalian carnivores found in the study area include snow leopard, common leopard (*Panthera pardus*), Himalayan lynx (*Lynx lynx*), Pallas’s cat (*Otocolobus manul*), leopard cat (*Prionailurus bengalensis*), grey wolf (*Canis lupus*), red fox (*Vulpes vulpes*), Asiatic jackal (*Canis aureus*), Himalayan brown bear (*Ursus arctos isabellinus*), Asiatic black bear (*Ursus thibetanus*), stone marten (*Martes foina*) and yellow-throated marten (*Martes flavigula*). Large-sized prey species include markhor (*Capra falconeri*), Himalayan ibex (*Capra siberica*), blue sheep (*Pseudois nayaur*), Ladakh urial (*Ovis vignei vignei*), Kashmir musk deer (*Moschus cupreus*), and Marco Polo sheep (*Ovis ammon polii*) (Ahmad et al. 2022).

**Figure 1:**
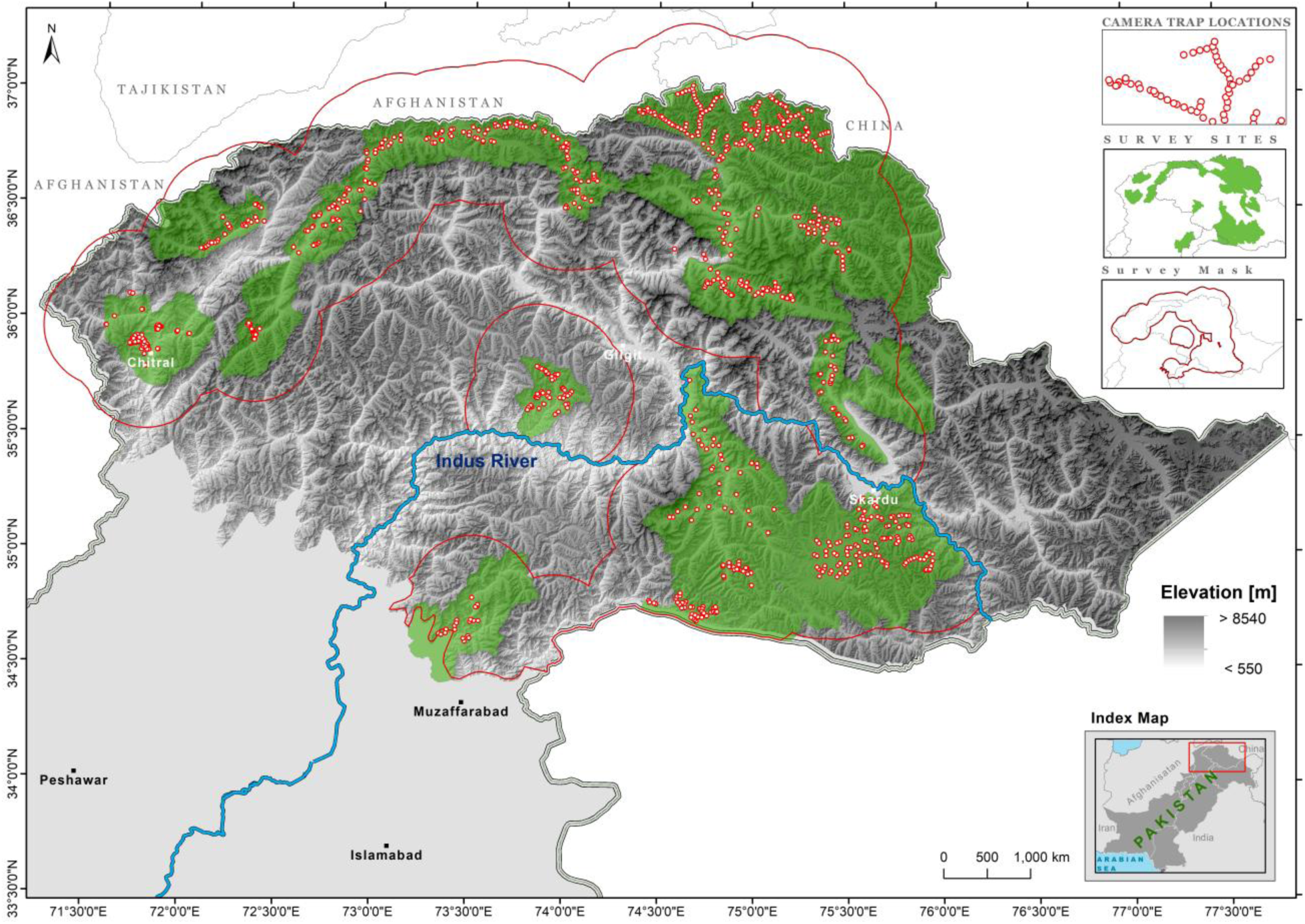
Spatial coverage of camera trapping carried out in Northern Pakistan from 2010-2019 to monitor snow leopards. The survey mask, defined as any potential habitat within 30 km of a camera station, is shown in red (60,684 km^2^). The Indus River acts as a barrier to snow leopard movement and is shown as a blue line. Camera locations are shown in red.

### Data Collection

Camera-trapping surveys are a widespread method for estimating population size and distribution, especially when individuals can be identified by, e.g., distinct pelage patterns, creating detection records of individual animals at each camera site (Rovero and Zimmermann 2016). A camera trapping study was conducted from 2010-2019 in 19 valleys across the snow leopard range in Pakistan (Figure 1, **Table 1**). A total of 828 cameras were installed and remained in the field for a total of 26,540 trap days (with an average of 32 days for each camera). Three different camera types were used: Reconyx, Suntek, and CamTrekker. Cameras were mostly installed on tracks, and the site for camera installation was selected based on animal signs such as pug marks, scrapes, evidence of scent marking, scat, or sightings of prey species. The camera was fixed on a metal pole with the sensor about 40-50 cm above the ground. Camera sensors were directed either north or south to avoid the rising or setting sun, which can cause false triggers (Ahmad et al. 2016). All the vegetation in front of the camera was removed, similarly to reduce false triggers. A lure (fish oil and castor oil) was spread on a stone in front of the majority of the cameras in an attempt to increase capture probabilities (Bischof et al., 2014).

**Table 1:** Summary statistics for the 19 camera trap surveys conducted to assess snow leopard density in Pakistan. Area of camera MCP = area of the minimum convex polygon bounding camera locations. The final column provides estimated abundances over the SCR mask associated with each survey, with 95% confidence intervals in parentheses, obtained from the model with the highest AIC (see Table 3). Estimated abundance for the entire survey region is given in Table 3. Administrative units include: KP = Khyber Pakhtunkhwa; GB = Gilgit-Baltistan, and AJK= Azad Jammu and Kashmir.

| Region | Start date | End date | Cameras | Mean effort per camera (days) | Area of camera MCP (km<sup>2</sup>) | Area of mask (km<sup>2</sup>) | Min distance between cameras | Cameras separated by < 750m | Detections | Animals | Cameras with detections | Estimated abundance |
| --- | --- | --- | --- | --- | --- | --- | --- | --- | --- | --- | --- | --- |
| Laspur Valley, KP | 2010/05/21 | 2010/06/23 | 20 | 32 | 21 | 3,455 | 573 | 20 | 1 | 1 | 1 | 4.3 (2.1-11) |
| Khunjerab National Park, GB | 2010/12/04 | 2011/01/03 | 10 | 29 | 92 | 4,811 | 1,908 | 1 | 12 | 8 | 4 | 22 (15.3-34.7) |
| Khunjerab National Park, GB | 2011/09/25 | 2011/11/16 | 86 | 11 | 1,036 | 7,722 | 1,363 | 2 | 24 | 15 | 14 | 31.9 (24-46.8) |
| Shimshal Valley, GB | 2011/10/01 | 2011/11/15 | 36 | 11 | 362 | 5,654 | 1,778 | 2 | 7 | 4 | 6 | 21.1 (13-36.3) |
| Tooshi-Chitral Gol National Park, KP | 2011/12/09 | 2012/01/26 | 22 | 41 | 698 | 6,636 | 924 | 7 | 0 | 0 | 0 | 1.8 (0.4-7.4) |
| Broghil National Park, KP and Qurumber National Park, GB | 2012/06/18 | 2012/07/30 | 80 | 15 | 1,794 | 9,891 | 1,373 | 3 | 1 | 1 | 1 | 11.5 (5.9-23.4) |
| Deosai National Park, GB | 2012/09/23 | 2012/11/09 | 116 | 16 | 1,530 | 6,980 | 1,505 | 2 | 0 | 0 | 0 | 1.5 (0.3-7.7) |
| Yarkhun Valley, KP | 2012/12/15 | 2013/01/23 | 58 | 32 | 849 | 8,292 | 1,569 | 6 | 0 | 0 | 0 | 5.5 (2.1-14.2) |
| Misgar Valley, GB | 2013/05/11 | 2013/06/14 | 59 | 31 | 691 | 6,789 | 1,579 | 2 | 24 | 5 | 13 | 17.7 (11.3-30.8) |
| Astore Valley, GB | 2013/06/23 | 2013/07/07 | 25 | 10 | 110 | 3,993 | 840 | 11 | 0 | 0 | 0 | 0.4 (0-3.9) |
| Musk Deer National Park, AJK | 2014/05/11 | 2014/06/26 | 40 | 27 | 227 | 3,338 | 1,083 | 3 | 1 | 1 | 1 | 1.2 (1-4.1) |
| Khanbari Valley, GB | 2014/12/04 | 2015/01/11 | 48 | 31 | 336 | 4,569 | 986 | 11 | 0 | 0 | 0 | 0.1 (0-2.4) |
| Terich Valley, KP | 2015/11/14 | 2015/12/30 | 28 | 40 | 253 | 5,383 | 1,375 | 2 | 1 | 1 | 1 | 5.7 (2.8-13.3) |
| Hopper_Hisper Valleys, GB | 2016/03/03 | 2016/05/07 | 38 | 53 | 258 | 5,781 | 1,408 | 2 | 34 | 4 | 10 | 9.5 (6.2-17.9) |
| Basha Valley, GB | 2017/05/20 | 2017/07/19 | 30 | 53 | 528 | 5,619 | 1,447 | 3 | 3 | 3 | 3 | 8 (4.9-16) |
| Hunza-Nagar Valleys, GB | 2018/04/11 | 2018/07/01 | 44 | 62 | 2,030 | 10,034 | 3,109 | 0 | 21 | 9 | 10 | 19.2 (13.9-30.6) |
| Kaghan Valley, KP | 2018/09/16 | 2018/10/06 | 20 | 16 | 590 | 4,233 | 3,410 | 0 | 0 | 0 | 0 | 0.1 (0-2.2) |
| Astore Valley, GB | 2018/11/20 | 2019/04/29 | 38 | 123 | 2,120 | 7,847 | 4,542 | 0 | 4 | 1 | 1 | 2.4 (1.3-7.7) |
| Chitral Gol National Park, KP | 2019/06/17 | 2019/08/06 | 30 | 46 | 46 | 3,686 | 911 | 9 | 0 | 0 | 0 | 0.2 (0-3.2) |
| **All** | **2010/05/21** | **2019/08/06** | **828** | **32** | **13,568** | **114,712** | **573** | **86** | **133** | **53** | **65** |  |

All of the cameras were set to take three consecutive photos (1 s apart) each time they were triggered. We also noted the habitat, substrate, topography, terrain, and altitude at each camera location.

Individual snow leopards were identified from photographs based on their distinct pelage patterns following Sharma et al. (2014) and Alexander et al. (2015) (Figure 2). To avoid identification errors, we adopted a two-step individual identification process. First, two teams independently checked all snow leopard photos and assigned individual IDs based on their unique coat patterns. In the second confirmation step, individual capture histories were compared, and any inconsistencies were resolved through discussion (Nawaz et al. 2021). We excluded blurred pictures in which individual identification was impossible.

**Figure 2:**
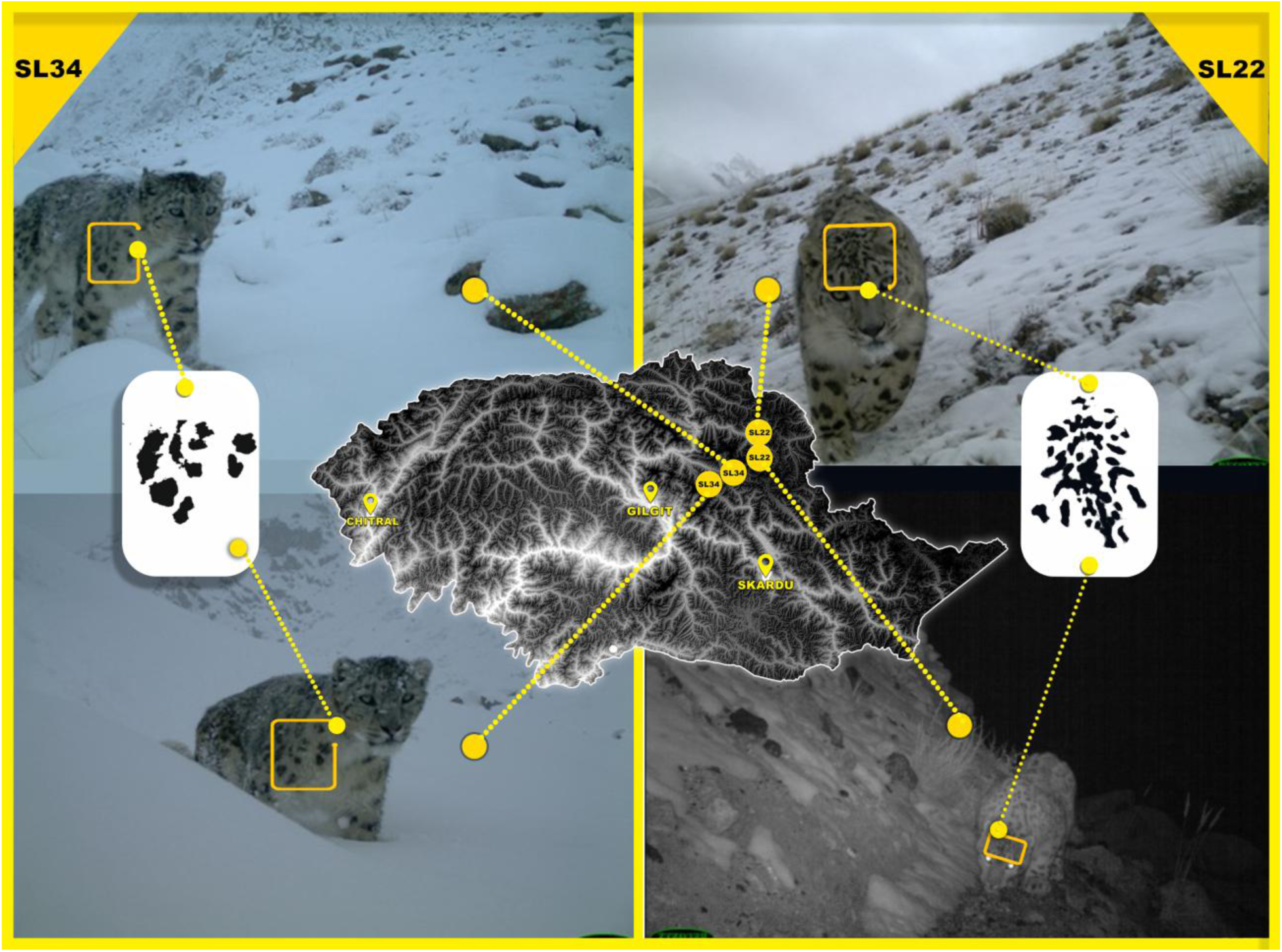
Photographic identification of individual snow leopards based on unique coat patterns. Individual named SL34 (Left top and bottom, captured from Hopper Valley) was identified from coat patterns on the shoulders. Individuals named SL22 (Right top and bottom, captured from Shimshal Valley), was identified from coat patterns on the forehead.

### Data Analysis

Estimates of snow leopard density were produced using maximum likelihood SCR methods implemented in the secr v4.6.5 package (Efford 2024) in R v4.3 (R Core Team 2023). SCR models (Borchers and Efford 2008, Royle and Young 2008) have been widely applied to estimate animal abundance from a variety of data types, including camera trap surveys, for many species. The methods integrate a spatial model that quantifies the density of latent animal activity centers across the survey region with an encounter model that assumes that the expected detection rate or probability decreases as the distance between the detector and the animal activity center increases.

We fitted SCR models to data from 19 independent surveys conducted between May 2010 and August 2019. Some of these surveys did not follow current best practice guidelines on camera spacing for snow leopard density with SCR methods, for example, due to: limited resources and experience, competing survey objectives, or because surveys pre-dated the development of the guidelines (Sharma et al., 2020; Dupont et al., 2021). In particular, the areas covered by camera trap arrays in nine of 19 surveys are smaller than 500 km^2^, the approximate typical size of a snow leopard home range (Johansson et al., 2016, **Table 1**). Surveys covering less than a home range run an increased risk of underestimating the spatial scale parameter of the encounter rate function (σ) and overestimating density (Harmsen et al., 2020; Sollmann et al., 2012; Nawaz et al., 2021). To check the sensitivity of our results to the inclusion of particular surveys, we ranked surveys by size (measured by the maximum distance between a camera pair) and removed surveys in ascending order of their sizes, i.e., from smallest to largest, fitting models to the remaining surveys. To assess whether any single survey exerted a disproportionate influence on results, we removed each survey one at a time and fitted models to the remaining 18 surveys. As results were broadly insensitive to survey selection, we present these results in **Appendix A**. Across all surveys, the average nearest-neighbor distance between cameras was 1,633 m, although this distance varied substantially between surveys and was less than 1 km in five of the 19 surveys (**Table 1**).

The 19 surveys analyzed here had either large distances between them (so that animals detected in one survey had no chance of being detected at another survey) or substantial time between them (so that the standard SCR assumptions of a closed population – no births, deaths, immigration or emigration over the survey period – was unlikely to hold). Due to many regions being sampled once, and sample sizes within each survey being small, open-population SCR methods were not feasible. We therefore treated each survey as an independent session and fitted multi-session SCR models to the data (Williams et al. 2021). Due to small numbers of detections in many of the surveys, and because we expected spatial covariates to operate in the same way throughout the broader survey region, model parameters were constrained to be equal over sessions. The resulting density must therefore be interpreted as an expected average density over the period of the whole study, rather than a snapshot of a single point in time.

Parameter estimation by maximum likelihood SCR methods requires numerical integration of the likelihood function, which is achieved by defining a fine mesh of points called the habitat mask at which the likelihood can be evaluated. This mask has the effect of discretizing continuous space into grid cells centered on the mesh points. The extent of the mask is chosen so that animals with activity centers beyond the mask have a near-zero chance of detection. We used the union of buffered polygons up to 30 km from any detector to define the mask area for each session (Figure 1), with a 2 km spacing between mask points, well within conventional guidelines (Sharma et al. 2020). Any mask points above 5500 m elevation were removed as unsuitable habitats for snow leopard activity centers. This threshold was based on our knowledge of snow leopards in the region and previous studies (Nawaz et al. 2021). In addition, the Indus River forms a barrier that we assumed to be rarely crossed by snow leopards (blue dashed line, Figure 1). Any mask points falling on a different side of the river from where the cameras were placed were also removed.

We fit a number of models that allowed snow leopard density and encounter rates to vary as a function of environmental covariates. Density was modeled as a function of elevation, slope, terrain roughness, normalized difference vegetation index (NDVI), population density, settlement density, and road density. All potential density covariates were measured on continuous scales and were scaled to have a mean of zero and unit variance. Baseline encounter rates (parameter λ_0_) were modelled as constant or a function of: camera type (Reconyx, Suntek, or Camtrekker); habitat at the camera location (barren, forest, pasture, or scrub); terrain at the camera location (base of the cliff, draw, plateau, ridge, saddle or valley), and a binary indicator of lure use. Varying survey effort (i.e. recording duration) at different cameras was accounted for by allowing detection rates to depend on the duration of exposure to detectors or “usage”. Model selection was based on Akaike’s Information Criterion (AIC). Following model selection, country-specific abundance and density estimates were produced by using fitted models to make predictions across the entirety of the snow leopard’s range in Pakistan (Figure 1, purple boundary).

## Results

Of the 828 cameras deployed over 26,540 trap days, 65 camera traps recorded 4,712 snow leopard images, with a mean detection rate of 0.17 per day. Out of 142 detection events, nine detections were included with photos (299) unsuitable for individual snow leopard identification due to improper camera trap placement angles and blurriness. The remaining 4,363 images were used to identify 53 individual snow leopards (80 recaptures, **Table 1**). About half of the unique individuals (53%) were detected only once, while two individuals were detected 10 and 17 times, respectively, accounting for one-quarter of all recaptures (**Table 2**). The average recapture frequency was 2.28 times, excluding animals detected only once.

**Table 2:** Frequencies of detection. The table shows the number of individual snow leopards (row “Individuals”) detected different numbers of times (row “Detections”).

|  |  |  |  |  |  |  |  |  |  |
| --- | --- | --- | --- | --- | --- | --- | --- | --- | --- |
| **Detections** | 1 | 2 | 3 | 4 | 6 | 7 | 8 | 10 | 17 |
| **Individuals** | 28 | 9 | 4 | 6 | 2 | 1 | 1 | 1 | 1 |

Model selection was in favor of a model with positive density associated with elevation and higher baseline encounter rates with Reconyx cameras than other types of cameras (AIC weight 60%; **Table 3**). The only other model with substantial support (AIC weight 27%) had density negatively associated with human population density with the same encounter rate function. No other model gained substantial support (all other AIC weights <8%), although all density covariate models outperformed the null model with constant density. All covariate relationships were in the expected direction (density positively associated with altitude, slope, and terrain ruggedness, and negatively associated with NDVI, population density, settlement density, and road density (Figure 3). Confidence intervals for all covariate effects did not contain zero, except for road density, which was extremely close (95% CI-1.26-0.01).

**Figure 3:**
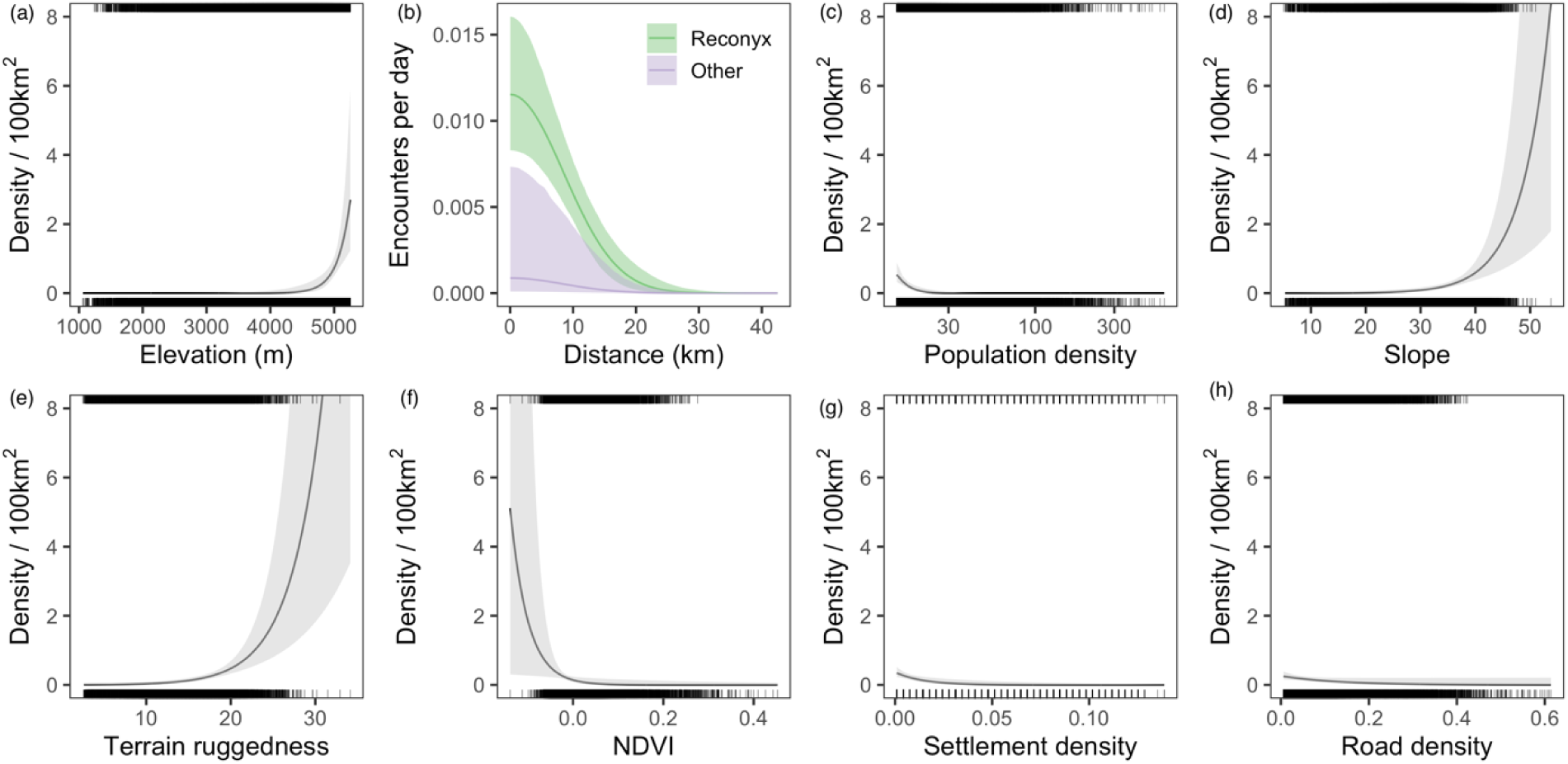
Changes in expected density of snow leopard activity centers (a, c-j) and expected encounter rate (b) as a function of biogeographical covariates. Panel (a) and (b) are obtained from the model with the highest AIC (Table 3, row 1). Panels (c) to (j) are obtained from different models each fitting the displayed covariate only. The solid line indicates the expected density or encounter rate, while the grey envelope shows 95% confidence intervals around that expected value. Rug plots at the top and bottom of plots show the distribution of covariate values in the mask region used to fit the model (top) and the full survey region used for prediction (bottom).

**Figure 4:**
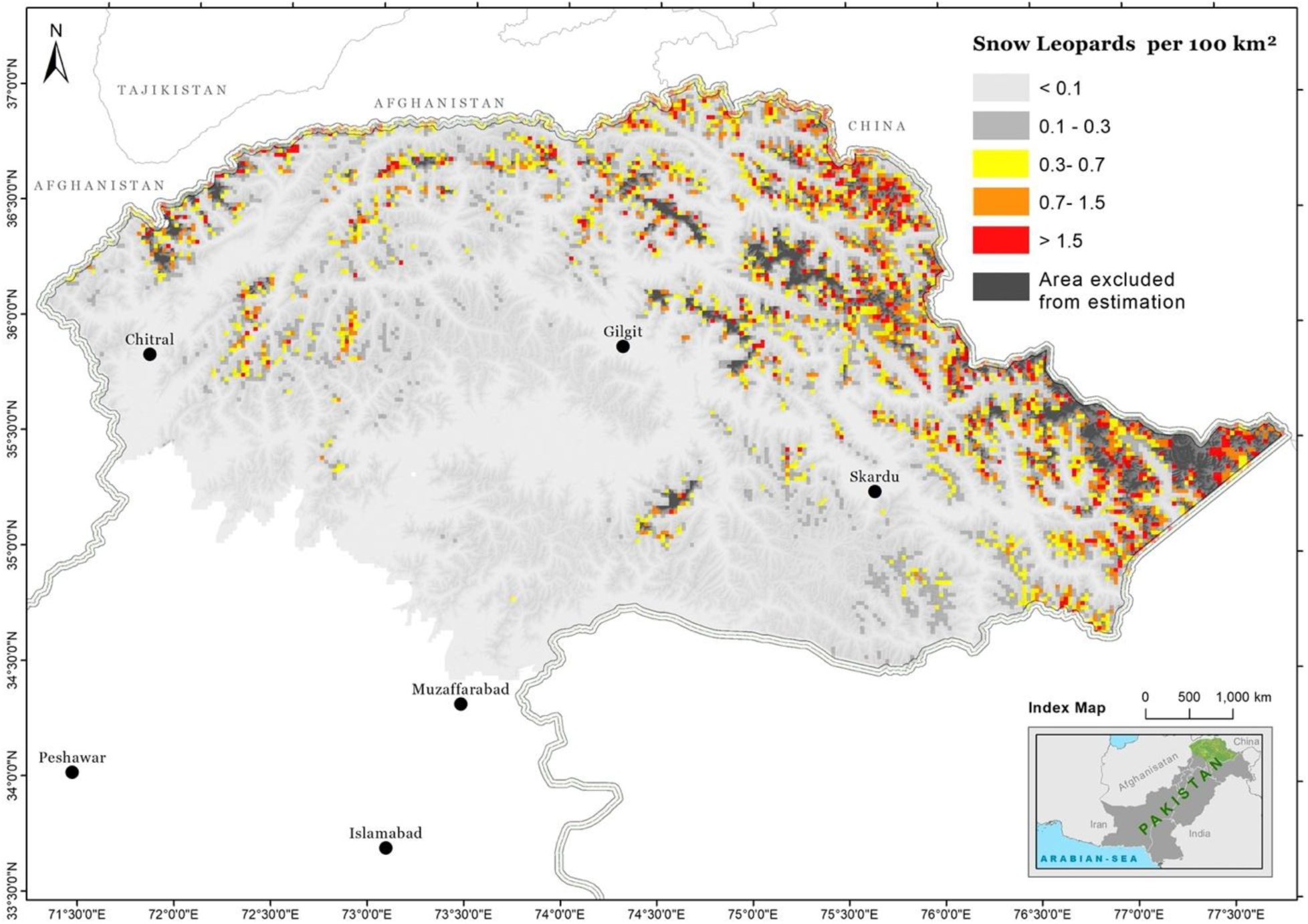
Spatial pattern of snow leopard density in Pakistan, derived from SCR model using camera trap data collected from 2010-2019.

**Table 3:** Model selection (AIC and AIC weights) for fitted models, with density and abundance estimates from each model to show the sensitivity of these estimates to the fitted model. Two sets of abundance and density estimates are given; one (labelled “Mask”) for the area covered by those parts of the survey mesh areas that fall within the survey region (Figure 2, areas within both red and purple borders); and one (labelled “Pakistan”) extrapolating to all snow leopard habitat across Pakistan (Figure 2, area within blue borders).

| Density Covariate | Detection Covariate | AIC | AIC Weight (%) | $\hat{N}$ (mask) | $\hat{D}$ (mask) | CV% | $\hat{N}$ (Pakistan) | $\hat{D}$ (Pakistan) |
| --- | --- | --- | --- | --- | --- | --- | --- | --- |
| Elevation | Camera type | 893 | 60 | 74 (51-106) | 0.12 (0.08-0.18) | 16 | 127 (88-182) | 0.13 (0.09-0.19) |
| Pop. Density | Camera type | 895 | 27 | 92 (62-134) | 0.15 (0.1-0.22) | 16 | 140 (96-204) | 0.14 (0.1-0.21) |
| Elevation | Terrain | 897 | 7 | 71 (49-102) | 0.12 (0.08-0.17) | 17 | 122 (85-175) | 0.12 (0.09-0.18) |
| Elevation | Habitat | 899 | 3 | 71 (49-104) | 0.12 (0.08-0.17) | 16 | 122 (85-177) | 0.12 (0.09-0.18) |
| Elevation | None | 900 | 2 | 71 (49-103) | 0.12 (0.08-0.17) | 17 | 123 (85-177) | 0.13 (0.09-0.18) |
| Pop. Density | None | 901 | 1 | 89 (61-131) | 0.15 (0.1-0.22) | 19 | 137 (94-200) | 0.14 (0.1-0.2) |
| Elevation | Lure used | 902 | 1 | 71 (49-104) | 0.12 (0.08-0.17) | 20 | 123 (85-177) | 0.13 (0.09-0.18) |
| Settlement Density | None | 920 | 0 | 115 (82-161) | 0.19 (0.13-0.26) | 17 | 201 (143-282) | 0.21 (0.15-0.29) |
| Slope | None | 921 | 0 | 100 (72-137) | 0.16 (0.12-0.23) | 16 | 169 (123-232) | 0.17 (0.13-0.24) |
| Terrain ruggedness | None | 922 | 0 | 100 (73-138) | 0.17 (0.12-0.23) | 16 | 169 (123-232) | 0.17 (0.13-0.24) |
| NDVI | None | 930 | 0 | 108 (77-151) | 0.18 (0.13-0.25) | 18 | 167 (119-234) | 0.17 (0.12-0.24) |
| Road Density | None | 934 | 0 | 102 (72-144) | 0.17 (0.12-0.24) | 18 | 166 (118-234) | 0.17 (0.12-0.24) |
| None | None | 937 | 0 | 103 (74-143) | 0.17 (0.12-0.24) | 17 | 166 (119-231) | 0.17 (0.12-0.24) |

The estimated density of snow leopards within the survey region was 0.12 animals per 100 km^2^ (95% CI 0.08–0.18; **Table 3**). Multiplying this density by the area of the survey mask region falling within Pakistan’s borders (60,684 km^2^), the estimated animal abundance is 74 (95% CI 51– 106). The precision of this abundance estimate is relatively high (CV = 16%) because the number of individuals and the number of recaptures made across all surveys were fairly high. Extrapolating to all potential habitats within Pakistan (98,284 km^2^) gives an estimated animal abundance of 127 (95% CI 88–182), equivalent to a mean density of 0.13 animals per 100 km^2^ (95% CI 0.09-0.19). The Karakoram range hosts the largest population (103 ± 19.5 snow leopards, average density 0.229 animals per 100 km^2^), followed by Hindukush (17.4 ± 3.66 snow leopards, average density 0.062), and Himalayas (6 ± 2.9 snow leopards, average density 0.024).

Snow leopard density varied significantly across the landscape, ranging from less than 0.1 to over 1.5 snow leopards per 100 km^2^, with the highest densities occurring towards high-elevation areas in northern Pakistan (Figure 3a). Density estimates were positively influenced by slope and ruggedness, while NDVI had a negative effect. Human population and infrastructure (Figure 3c, g, h) did not appear to affect the density pattern of snow leopards.

The scale parameter σ indicating how encounter rates decrease with distance from the activity center was estimated to be 8,498 m (95% CI 6,933–10,416), towards the higher end of what has been observed in other snow leopard surveys (Bian et al., 2023; Oberosler et al., 2022; Suryawanshi et al., 2021; Zhang et al., 2024). The baseline encounter rate λ_0_(the expected encounter rate for an animal whose activity center is at a camera) was estimated to be 0.011 encounters per day (95% CI 0.008–0.016) at Reconyx cameras and 0.0008 encounters per day (95% CI 0.0001–0.007) at other cameras (Figure 3b).

## Discussion

### Establishing the Baseline

The snow leopard is among the least-studied large carnivore species in Pakistan, with no reliable, empirically derived estimates of population size or spatial density patterns available. Accurate population estimates are crucial for conservation planning, as they enable management agencies to devise evidence-based policies to protect rare and threatened species (Gerber et al. 2014). However, such data remain scarce across most of the snow leopard’s range, including Pakistan (Snow Leopard Network, 2021). This study provides the first-ever robust, nationwide population estimate of snow leopards in Pakistan, derived from systematic camera-trapping methods.

The current study is significant as it establishes a benchmark for population, employing scientifically validated methods such as SCR and extensive sampling. The study covers 39% of snow leopard habitat, representing both high- and low-quality areas in Pakistan (Hameed et al., 2020). The SCR predictions were consistent with the prevailing understanding of density variation in the region. For example, the model accurately identifies high-density areas such as Gojal, Hopper-Hisper, Qurumber, Hushey, and Thallay valleys, which are characterized by suitable habitats both for snow leopards (Hameed et al. 2020) and their prey (Ali et al. 2021). Notably, the model predicted higher densities in good quality habitats ( like Hushey and Thallay in Baltistan, Hameed et al., 2020), which were not even covered in the camera trapping effort. These findings underscore the model’s robustness in capturing spatial density patterns across the landscape.

The predicted high snow leopard density is predominantly located in designated protected areas (PAs), including national parks (KNP, QNP, BNP, and CKNP) and community-controlled hunting areas, such as Passu, Khyber, KVO, Gulmit, Ghulkin, and Hussaini valleys. This indicates the important role of the PAs network in maintaining threatened species (Wen et al. 2022). These PAs provide a suitable habitat (Din et al. 2022b) and support a healthy population of large-sized prey species such as ibex and blue sheep (Ahmad et al. 2022), which is a pivotal factor of snow leopard density (Johansson et al. 2016). Ibex in northern Pakistan is an important source of food for snow leopards, and its populations alone could support 97 to 135 snow leopards (Ahmad et al., 2022). Most high-density ibex populations are confined to protected areas (Ahmad et al. 2022), reinforcing the correlation between snow leopard density and prey availability.

Reported densities for snow leopards exhibit considerable variability across the range, attributed to ecological, anthropogenic, and methodological factors (Nawaz et al., 2021; Pal et al., 2022; Suryawanshi et al., 2021). Density estimates have been documented to vary from as low as 0.12 individuals per 100 km² in India (Suryawanshi et al., 2021) to a maximum of 5.32 individuals per 100 km² in China (Alexander et al. 2015). For different countries, known density estimates per 100 km² in ascending order are as follows: 0.57 to 0.95 individuals in Tajikistan (Kachel et al. 2017); 0.66 to 4.6 in Nepal (Chetri et al. 2019, Khanal et al. 2020); 0.31 to 1.5 in Mongolia (Sharma et al. 2014, Bayandonoi et al. 2021, Oberosler et al. 2022); 0.12 to 2.15 in India (Sharma et al. 2021, Suryawanshi et al. 2021b, Pal et al. 2022); 0.90 in China (Alexander et al. 2015, 2016, Bian et al. 2023, Zhang et al. 2024, Li et al. 2025), and 1.2 to 1.48 in Bhutan (NCD 2023). In contrast, the current study estimates a mean snow leopard density of 0.13 animals per 100 km^2^ (95% CI 0.09-0.19), which is lower than most of the estimates reported for other range countries. However, direct comparisons among these studies are challenging due to substantial variability in their spatial coverage. Unlike the present study and that conducted in Bhutan (NCD 2023), Mongolia (Bayandonoi et al. 2021), and China (Li et al. 2025), which provide density estimates at a large spatial scale, most other studies focus on smaller areas, ranging from 375 km² to 5,484 km². Such localized studies are often not representative of broader spatial patterns across the species’ range, as they frequently target high-quality habitats (Suryawanshi et al., 2021). For instance, in Pakistan, certain patches have densities exceeding 1.5 individuals per 100 km², reflecting localized variation within the overall range.

Past subjective estimates of snow leopard populations in Pakistan, ranging from 250 to 400 individuals (Schaller 1976, Malik 1997, Hussain 2003), relied on indirect methods (sign surveys, habitat-based prediction, interviews, and expert judgments), often extrapolated from limited habitat surveys. In contrast, the current estimate of 127 individuals (95% CI 88-188), derived from true detections captured from 39% of the species’ range, represents a baseline correction rather than a population decline. The high estimated sigma value (8.5 km) indicates large-scale movements of individuals, further explaining potential overestimations in previous studies.

### A Decade of Lessons: Challenges and Innovations in Camera Trapping in Snow Leopard Habitat

Camera trapping in Pakistan represents a comprehensive endeavor, spanning over a decade and covering the vast span of the snow leopard’s habitat, including difficult terrain and extreme conditions. This undertaking commenced at a time when guidelines for camera trapping, specifically for population studies in snow leopard habitats, were nonexistent (Nawaz et al. 2021). Consequently, our efforts have involved considerable experimentation and course corrections, which have subsequently improved the efficacy of more recent surveys. We share our experiences, focusing on mistakes and adaptations, with the hope of guiding other researchers working in similar ecological systems.

We identified four critical factors that influence the success of camera trapping: season, study site size, camera configuration, and duration of sampling. Summer is a season with better accessibility in snow leopard terrain, however, wildlife detections are significantly reduced due to higher human and livestock movement during this season. For instance, our surveys in the Astore Valley in 2013, carried out in summer, yielded overall limited wildlife detections, with no snow leopard captures but frequent livestock images. In contrast, winter, though harsh and logistically challenging, offers higher detection rates at lower elevations.

Our team was less thoughtful about the size of camera trap grids during early surveys. For example, the survey area in the Laspur camera trapping of 2010 was smaller than the home range of a single snow leopard, violating recommendations for SCR sampling design (Sollmann et al. 2012). Sampling small areas, combined with a bias toward high-quality habitats, often leads to density overestimation by up to five times (Suryawanshi et al. 2019). Additionally, camera configurations remained inconsistent. In some valleys, cameras were placed too close together (<1 km apart, e.g., Hopper Valley in 2016), resulting in inflated density estimates (Nawaz et al. 2021).

Spatial coverage is a key consideration in ecological sampling, which can be achieved through methods such as systematic sampling, random sampling, or cluster sampling (Williams and Brown 2019, Nawaz et al. 2021). However, implementing these approaches in the rugged terrain of snow leopard habitats is extremely challenging. For example, our initial attempt to install cameras based on a 5 x 5 km sampling grid (e.g., Broghil NP in 2012) encountered multiple difficulties. Several grids were inaccessible due to rugged terrain, river crossings, and melting glaciers, resulting in incomplete sampling and compromising analytical accuracy.

To address these issues, we adopted a watershed-based approach. This approach involves dividing the study area into sampling units along the natural hydrological watersheds. This approach proved effective in several surveys (e.g., Musk Deer NP, Khanbari Valley in 2014), as it allowed for accessing all sampling units. This was because accessibility in highly rugged terrains typically follows river or stream basins. Although high-elevation and steeper areas remained inaccessible in certain units, animals were still detected as they moved along watershed courses. For example, in the Misgar study (**Table 1**), several recaptures were recorded across the elevation gradient of a watershed. Duration of sampling is another important consideration in population sampling (Sollmann et al. 2012). In some of our earlier studies, cameras were placed for two weeks (e.g., the 2011 studies in KNP and Shimshal), driven by the desire to cover larger areas with limited camera resources. However, subsequent analyses (e.g., Bischof et al., 2014) highlighted that a minimum of 40 days is required to optimize snow leopard detections.

Despite the issues identified here, sensitivity analyses (**Appendix 1**) indicated that abundance estimates reported in this study were broadly robust. This robustness is likely due to the pooling of data from several studies and the coverage of a vast spatial scale, which includes both good and poor-quality habitats.

## Conclusion and Recommendations

This study provides the first-ever robust, nationwide population estimate of snow leopards in Pakistan, derived from systematic camera-trapping methods. We estimate that up to 180 snow leopards may thrive in the territory of Pakistan. However, it is important to note that the current population estimate reflects an average over a decade of camera-trapping efforts, rather than a snapshot in a particular year. To validate these estimates and address temporal variability, future studies should integrate independent methods, such as non-invasive genetic analyses.

Despite its limitations, this study provides the first reliable population baseline for snow leopards in Pakistan, offering a foundation for monitoring future trends and assessing the effectiveness of ongoing conservation measures. Given the small population size, dispersed distribution, emerging threats such as climate change, and significant ecological role, the snow leopard should remain a high-priority species for conservation in High Asia. Dedicated efforts are essential to ensure the long-term viability of this iconic predator and the ecosystems it inhabits.

Wildlife biologists often initiate camera-trapping studies with the excitement of capturing snow leopard images but without sufficient attention to sampling design, resulting in data that cannot support robust population inferences (Suryawanshi et al. 2019). A predominant focus on high-quality habitats or the most promising detection sites can lead to biased population estimates (Sharma et al. 2018). Based on recent SCR sampling recommendations (e.g., Dupont et al., 2021; Sollmann et al., 2012; Tobler et al., 2013) and our field experiences, we offer guidance for effective camera trapping in snow leopard habitats. Seasonal considerations are critical, and the optimal period for camera trapping in snow leopard habitats is between October and March (autumn, winter, and early spring) when human and livestock interference is minimal. For reliable estimates, the study area should cover at least 481 km² (Suryawanshi et al. 2019), and larger spatial coverage affords more accurate estimates (Nawaz et al., 2021). Sampling should include both high- and low-quality habitats to ensure realistic and unbiased numbers (Suryawanshi et al. 2019, Durbach et al. 2021). Utilizing design tools and recommendations developed by Dupont et al. (2021) and Durbach et al. (2021) can help address the challenges in sampling imposed by the difficult terrain of High Asia. Cameras should be installed for a minimum of 40 days to optimize capture and recapture for snow leopards (Bischof et al., 2014). Additionally, the use of high-quality, reliable cameras is critical, as our analyses indicate a strong impact of camera type on detection success.

These recommendations, coupled with systematic and adaptive methodologies, will contribute to more accurate and effective population monitoring of snow leopards and other carnivores in the region.

## Supporting information

Appendix 1, sensitivity analysis

## Acknowledgments

We are grateful to the Snow Leopard Trust, GEF, Whitley Fund for Nature, Norwegian Research Council, and Panthera for their financial support, and to the Snow Leopard Foundation for field facilitation and logistics. We also appreciate the assistance provided by the staff of the Park and Wildlife Department of GB, the Fisheries and Wildlife Department of AJK, the KP Wildlife Department, and the District Administrations during our fieldwork. Additionally, several individuals from local communities, wildlife staff, and students supported data collection, particularly Ejaz Ur Rehman, Muhammad Asif, Nazakat Din, Fathul Bari, Muhammad Shakil, Barkat Ullah Khan, Siraj Khan, and Mohammad Younas. Doost Ali Nawaz helped with preparing GIS maps. Dr. Oliver Wearn reviewed the initial draft and provided helpful comments. We thank them all.

## Data Availability

The corresponding author will provide the datasets used and/or analyzed during the current study on request.

## Author Contributions

**Conceptualization**: Muhammad Ali Nawaz

**Data curation**: Shoaib Hameed, Hussain Ali, Jaffar ud Din

**Formal analysis**: Ian Durbatch, Shoaib Hameed, Muhammad Ali Nawaz

**Funding acquisition**: Muhammad Ali Nawaz

**Investigation**: Hussain Ali, Shoaib Hameed, Jaffar ud Din, Mehmood Ghaznavi, Mohsin Farooque, Naeem Iftikhar, Muhammad Samar Hussain Khan

**Methodology**: Hussain Ali, Shoaib Hameed, Shakeel Ahmad, Muhammad Ali Nawaz

**Project administration**: Jaffar ud Din, Muhammad Ali Nawaz

**Supervision**: Muhammad Ali Nawaz

**Visualization**: Shoaib Hameed, Shakeel Ahmad, Hussain Ali

**Writing** – **original draft**: Muhammad Ali Nawaz, Shakeel Ahmad, Ian Durbatch

**Writing – review & editing**: Muhammad Ali Nawaz, Ian Durbatch

## References

Ahmad, S., Hameed, S., Ali, H., Khan, T. U., Mehmood, T. and Nawaz, M. A. 2016. Carnivores’ diversity and conflicts with humans in Musk Deer National Park, Azad Jammu and Kashmir, Pakistan. - Eur. J. Wildl. Res. 62: 565–576.

Ahmad, S., Ali, H., Asif, M., Khan, T., Din, N., Rehman, E. U., Hameed, S., Din, J. U. and Nawaz, M. A. 2022. Spatial density pattern of Himalayan Ibex (Capra sibirica) in Pakistan. - Glob. Ecol. Conserv. 39: e02288.

Ale, S. B. and Mishra, C. 2018. The snow leopard’s questionable comeback. - Science (80-.). in press.

Alexander, J. S., Gopalaswamy, A. M., Shi, K., Riordan, P. and Margalida, A. 2015. Face value: Towards robust estimates of snow leopard densities. - PLoS One in press.

Alexander, J. S., Zhang, C., Shi, K. and Riordan, P. 2016. A granular view of a snow leopard population using camera traps in Central China. - Biol. Conserv. in press.

Ali, S., Gao, J., Begum, F., Rasool, A., Ismail, M., Cai, Y., Ali, S. and Ali, S. 2017. Health assessment using aqua-quality indicators of alpine streams (Khunjerab National Park), Gilgit, Pakistan.: 4685–4698.

Ali, H., Din, J. U., Bosso, L., Hameed, S., Kabir, M., Younas, M. and Nawaz, M. A. 2021. Expanding or shrinking? range shifts in wild ungulates under climate change in Pamir-Karakoram Mountains, Pakistan. - PLoS One in press.

Balme, G., Rogan, M., Thomas, L., Pitman, R., Mann, G., Whittington-Jones, G., Midlane, N., Broodryk, M., Broodryk, K., Campbell, M., Alkema, M., Wright, D. and Hunter, L. 2019. Big cats at large: Density, structure, and spatio-temporal patterns of a leopard population free of anthropogenic mortality. - Popul. Ecol. in press.

Bayandonoi, G., Lkhagvajav, P., Alexander, J. S., Durbach, I., Borchers, D., Munkhtsog, B. and Sharma, K. 2021. Nationwide snow leopard population assessment of Mongolia: key findings. Summary report. Ulaan baatar, Mongolia.

Bian, X., Liang, X., Weckworth, B., Jyal, D., Hull, V. and Yang, L. 2023. Spatial density estimate of the snow leopard, Panthera uncia, in the Central Tibetan Plateau, China. - Integr. Zool. in press.

Bischof, R., Ali, H., Kabir, M., Hameed, S. and Nawaz, M. A. 2014a. Being the underdog: An elusive small carnivore uses space with prey and time without enemies. - J. Zool. in press.

Bischof, R., Hameed, S., Ali, H., Kabir, M., Younas, M., Shah, K. A., Din, J. U. and Nawaz, M. A. 2014b. Using time-to-event analysis to complement hierarchical methods when assessing determinants of photographic detectability during camera trapping. - Methods Ecol. Evol. 5: 44–53.

Borchers, D. L. and Efford, M. G. 2008. Spatially explicit maximum likelihood methods for capture-recapture studies. - Biometrics in press.

Chetri, M., Odden, M., Sharma, K. and Flagstad, Ø. 2019. Estimating snow leopard density using fecal DNA in a large landscape in north-central Nepal. - Glob. Ecol. Conserv. 17: e00548.

Crooks, K. R., Burdett, C. L., Theobald, D. M., Rondinini, C. and Boitani, L. 2011. Global patterns of fragmentation and connectivity of mammalian carnivore habitat. - Philos. Trans. R. Soc. B Biol. Sci. in press.

Darimont, C. T., Paquet, P. C., Treves, A., Artelle, K. A. and Chapron, G. 2018. Political populations of large carnivores. - Conserv. Biol. in press.

Din, J. U., Bari, F., Ali, H., Rehman, E. ur, Hasan Adli, D. S., Abdullah, N. A., Norma-Rashid, Y., Kabir, M., Hameed, S., Nawaz, D. A. and Nawaz, M. A. 2022a. Drivers of snow leopard poaching and trade in Pakistan and implications for management. - Nat. Conserv. in press.

Din, J. U., Hameed, S., Ali, H., Norma-Rashid, Y., Hasan Adli, D. S. and Nawaz, M. A. 2022b. On the snow leopard Trails: Occupancy pattern and implications for management in the Pamir. - Saudi J. Biol. Sci. in press.

Din, J. U., Hameed, S., Ali, H. and Nawaz, M. A. 2023. The current state of snow leopard conservation in Pakistan. - In: Snow Leopards. in press.

Dupont, G., Royle, J. A., Nawaz, M. A. and Sutherland, C. 2021. Optimal sampling design for spatial capture–recapture. - Ecology in press.

Durbach, I., Borchers, D., Sutherland, C. and Sharma, K. 2021. Fast, flexible alternatives to regular grid designs for spatial capture–recapture. - Methods Ecol. Evol. in press.

Efford, M. 2024. secr: Spatially explicit capture-recapture models. in press.

Elliot, N. B., Bett, A., Chege, M., Sankan, K., de Souza, N., Kariuki, L., Broekhuis, F., Omondi, P., Ngene, S. and Gopalaswamy, A. M. 2020. The importance of reliable monitoring methods for the management of small, isolated populations. - Conserv. Sci. Pract. in press.

Gerber, B. D., Ivan, J. S. and Burnham, K. P. 2014. Estimating the abundance of rare and elusive carnivores from photographic-sampling data when the population size is very small. - Popul. Ecol. in press.

Gopalaswamy, A. M., Ullas Karanth, K., Delampady, M. and Stenseth, N. C. 2019. How sampling-based overdispersion reveals India’s tiger monitoring orthodoxy. - Conserv. Sci. Pract. in press.

Hameed, S., ud Din, J., Ali, H., Kabir, M., Younas, M., Ur Rehman, E., Bari, F., Hao, W., Bischof, R. and Nawaz, M. A. 2020. Identifying priority landscapes for conservation of snow leopards in Pakistan. - PLoS One in press.

Harmsen, B. J., Foster, R. J. and Quigley, H. 2020. Spatially explicit capture recapture density estimates: Robustness, accuracy and precision in a long-term study of jaguars (Panthera onca). - PLoS One in press.

Hussain, S. 2003. The status of the snow leopard in Pakistan and its conflict with local farmers. - ORYX in press.

Jackson, R. M., Roe, J. D., Wangchuk, R. and Hunter, D. O. 2006. Estimating Snow Leopard Population Abundance Using Photography and Capture–Recapture Techniques. - Wildl. Soc. Bull. in press.

Janečka, J. E., Jackson, R., Yuquang, Z., Diqiang, L., Munkhtsog, B., Buckley-Beason, V. and Murphy, W. J. 2008. Population monitoring of snow leopards using noninvasive collection of scat samples: A pilot study. - Anim. Conserv. in press.

Janečka, J. E., Munkhtsog, B., Jackson, R. M., Naranbaatar, G., Mallon, D. P. and Murphy, W. J. 2011. Comparison of noninvasive genetic and camera-trapping techniques for surveying snow leopards. - J. Mammal. in press.

Janjua, S., Peters, J. L., Weckworth, B., Abbas, F. I., Bahn, V., Johansson, O. and Rooney, T. P. 2020. Improving our conservation genetic toolkit: ddRAD-seq for SNPs in snow leopards. - Conserv. Genet. Resour. 12: 257–261.

Johansson, Ö., Rauset, G. R., Samelius, G., McCarthy, T., Andrén, H., Tumursukh, L. and Mishra, C. 2016. Land sharing is essential for snow leopard conservation. - Biol. Conserv. in press.

Johansson, Ö., Samelius, G., Wikberg, E., Chapron, G., Mishra, C. and Low, M. 2020. Identification errors in camera-trap studies result in systematic population overestimation. - Sci. Rep. in press.

Jones, J. P. G., Andriamarovololona, M. M., Hockley, N., Gibbons, J. M. and Milner-Gulland, E. J. 2008. Testing the use of interviews as a tool for monitoring trends in the harvesting of wild species. - J. Appl. Ecol. in press.

Kachel, S. M., McCarthy, K. P., McCarthy, T. M. and Oshurmamadov, N. 2017. Investigating the potential impact of trophy hunting of wild ungulates on snow leopard Panthera uncia conservation in Tajikistan. - ORYX in press.

Khanal, G., Mishra, C. and Ramesh Suryawanshi, K. 2020. Relative influence of wild prey and livestock abundance on carnivore-caused livestock predation. - Ecol. Evol. in press.

Li, J., Weckworth, B. V., McCarthy, T. M., Liang, X., Liu, Y., Xing, R., Li, D., Zhang, Y., Xue, Y., Jackson, R., Xiao, L., Cheng, C., Li, S., Xu, F., Ma, M., Yang, X., Diao, K., Gao, Y., Song, D., Nowell, K., He, B., Li, Y., McCarthy, K., Paltsyn, M. Y., Sharma, K., Mishra, C., Schaller, G. B., Lu, Z. and Beissinger, S. R. 2020. Defining priorities for global snow leopard conservation landscapes. - Biol. Conserv. in press.

Li, X., Wei, C., Chen, X., Jia, D., Li, P., Liang, S., Jikmed, A., Gao, Y., Zhao, X., Chu, M. and Sharma, K. 2025. First large - scale assessment of snow leopard population in China using existing data from multiple organizations. - Biodivers. Conserv. in press.

Linden, D. W., Green, D. S., Chelysheva, E. V., Mandere, S. M. and Dloniak, S. M. 2020. Challenges and opportunities in population monitoring of cheetahs. - Popul. Ecol. in press.

Malik, M. M. 1997. Status and Conservation of Snow Leopard in Pakistan. - Proc. Eighth Int. Snow Leopard Symp.: 11–20.

McCarthy, T. M. and Chapron, G. 2014. Snow Leopard Survival Strategy: Revised 2014 Version. - Snow Leopard Netw. in press.

McCarthy, T., Mallon, D., Jackson, R., Zahler, P. and McCarthy, K. 2017. Panthera uncia. The IUCN Red List of Threatened Species 2017: e.T22732A50664030. 10.2305/IUCN.UK.2017-2.RLTS.T22732A50664030.en. (Accessed 3 June 2024).

Moqanaki, E. M., Jiménez, J., Bensch, S. and López-Bao, J. V. 2018. Counting bears in the Iranian Caucasus: Remarkable mismatch between scientifically-sound population estimates and perceptions. - Biol. Conserv. in press.

Nawaz, M. A., Khan, B. U., Mahmood, A., Younas, M., Din, J. and Sutherland, C. 2021. An empirical demonstration of the effect of study design on density estimations. - Sci. Rep.: 1– 10.

NCD 2023. Snow leopard status in Bhutan: National snow leopard survey report2022-2023. Nature Conservation Division, Department of Forests and Park Services,Ministry of Energy and Natural Resources, Royal Government of Bhutan, Thimphu, Bhutan (PDF) Snow Leopard Stat.

Oberosler, V., Tenan, S., Groff, C., Krofel, M., Augugliaro, C., Munkhtsog, B. and Rovero, F. 2022. First spatially-explicit density estimate for a snow leopard population in the Altai Mountains. - Biodivers. Conserv. in press.

Pal, R., Sutherland, C., Qureshi, Q. and Sathyakumar, S. 2022. Landscape connectivity and population density of snow leopards across a multi-use landscape in Western Himalaya. - Anim. Conserv. in press.

R Core Team 2023. R: A Language and Environment for Statistical Computing. R Foundation for Statistical Computing, Vienna, Austria. in press.

Rovero, F. and Zimmermann, F. 2016. Camera trapping for Wildlife Research.

Royle, J. A. and Young, K. V. 2008. A hierarchical model for spatial capture recapture data. - Ecology in press.

Royle, J. A., Chandler, R. B., Sollmann, R. and Gardner, B. 2014. Spatial Capture-Recapture. - Academic Press.

Sayed, A. H., Bailey, A. J., Iqbal, M. M., Khan, A. M., Tuesday, L. U., Intended, T., Determined, N., Climate, U. N., Conference, C., Conservation, G., Gilgit, J., Mics, E. C. O. N. O., Sterner, T., Park, A. L. and Vidal, J. 2013. World Wide Fund for Nature-Pakistan Climate Change in the Northern Areas Pakistan Impacts on glaciers, ecology and livelyhoods. - Prog. Hum. Geogr. in press.

Schaller, G. B. 1976. Mountain Mammals in Pakistan. - Oryx in press.

Sharief, A., Kumar, V., Singh, H., Mukherjee, T., Dutta, R., Joshi, B. D., Bhattacharjee, S., Ramesh, C., Chandra, K., Thakur, M. and Sharma, L. K. 2022. Landscape use and co-occurrence pattern of snow leopard (Panthera uncia) and its prey species in the fragile ecosystem of Spiti Valley, Himachal Pradesh. - PLoS One in press.

Sharma, K., Bayrakcismith, R., Tumursukh, L., Johansson, O., Sevger, P., McCarthy, T. and Mishra, C. 2014. Vigorous dynamics underlie a stable population of the endangered snow leopard Panthera uncia in Tost Mountains, South Gobi, Mongolia. - PLoS One in press.

Sharma, K., Borchers, D., Tumursukh, L., Lkhagvajav, P. and Mishra, C. 2018. Understanding snow leopard distribution and their spatial ecology through Spatial Capture Recapture Analysis. - Int. Stat. Ecol. Conf. 2018 St Andrews, UK

Sharma, K., Borchers, D., Mackenzie, D., Durbach, I., Sutherland, C., Nichols, J., Lovari, S., Zhi, L., Ale, S., Khan, A. A., Modaqiq, W., McCarthy, T., Alexander, J. S. and Mishra, C. 2020. PAWS Guidelines.: 1–62.

Sharma, R. K., Sharma, K., Borchers, D., Bhatnagar, Y. V., Suryawanshi, K. R. and Mishra, C. 2021. Spatial variation in population-density of snow leopards in a multiple use landscape in Spiti Valley, Trans-Himalaya. - PLoS One in press.

Shedayi, A. A., Xu, M., Hussain, F., Sadia, S., Naseer, I. and Bano, S. 2016. Threatened plant resources: Distribution and ecosystem services in the world’s high elevation park of the karakoram ranges. - Pakistan J. Bot. in press.

Shen, Q. 2020. The Effects of Climate Change on Snow Leopards at the Hengduan Mountain Region. - IOP Conf. Ser. Earth Environ. Sci. 552

Sollmann, R., Gardner, B. and Belant, J. L. 2012. How does spatial study design influence density estimates from spatial capture-recapture models? - PLoS One in press.

Suryawanshi, K. R., Khanyari, M., Sharma, K., Lkhagvajav, P. and Mishra, C. 2019. Sampling bias in snow leopard population estimation studies. - Popul. Ecol. in press.

Suryawanshi, K., Reddy, A., Sharma, M., Khanyari, M., Bijoor, A., Rathore, D., Jaggi, H., Khara, A., Malgaonkar, A., Ghoshal, A., Patel, J. and Mishra, C. 2021a. Estimating snow leopard and prey populations at large spatial scales. - Ecol. Solut. Evid. 2: e12115.

Suryawanshi, K., Reddy, A., Sharma, M., Khanyari, M., Bijoor, A., Rathore, D., Jaggi, H., Khara, A., Malgaonkar, A., Ghoshal, A., Patel, J. and Mishra, C. 2021b. Estimating snow leopard and prey populations at large spatial scales. - Ecol. Solut. Evid. in press.

Tobler, M. W., Carrillo-Percastegui, S. E., Zúñiga Hartley, A. and Powell, G. V. N. 2013. High jaguar densities and large population sizes in the core habitat of the southwestern Amazon. - Biol. Conserv. in press.

Trouwborst, A. 2015. Global large carnivore conservation and international law. - Biodivers. Conserv. in press.

Watts, S. M., McCarthy, T. M. and Namgail, T. 2019. Modelling potential habitat for snow leopards (Panthera uncia) in Ladakh, India. - PLoS One in press.

Weise, F. J., Vijay, V., Jacobson, A. P., Schoonover, R. F., Groom, R. J., Horgan, J., Keeping, D., Klein, R., Marnewick, K., Maude, G., Melzheimer, J., Mills, G., van der Merwe, V., van der Meer, E., van Vuuren, R. J., Wachter, B. and Pimm, S. L. 2017. The distribution and numbers of cheetah (Acinonyx jubatus) in southern Africa. - PeerJ in press.

Wen, D., Qi, J., Long, Z., Gu, J., Tian, Y., Roberts, N. J., Yang, E., Kong, W., Zhao, Y., Sun, Q. and Jiang, G. 2022. Conservation potentials and limitations of large carnivores in protected areas: A case study in Northeast China. - Conserv. Sci. Pract. in press.

Williams, B. K. and Brown, E. D. 2019. Sampling and analysis frameworks for inference in ecology. - Methods Ecol. Evol. in press.

Williams, K. S., Williams, S. T., Welch, R. J., Marneweck, C. J., Mann, G. K. H., Pitman, R. T., Whittington-Jones, G., Balme, G. A., Parker, D. M. and Hill, R. A. 2021. Assumptions about fence permeability influence density estimates for brown hyaenas across South Africa. - Sci. Rep. in press.

Zaman, M., Chen, Y., Jackson, R. and Hussain, S. 2024. Silent Signals in the Snow: Tracking the Spatio--Temporal Territorial Marking Behavior of Snow Leopards (Panthera uncia) in the Mountainous Region of Baltistan, Pakistan.: 1–16.

Zhang, C., Ma, T. and Ma, D. 2024. Status of the snow leopard Panthera uncia in the Qilian Mountains, Gansu Province, China. - ORYX in press.

