## Appendix 1, sensitivity analysis for "From Shadows to Data: A Robust Population Assessment of Snow Leopards in the Highland Crossroads"

**Appendix A: Sensitivity of estimates to survey inclusion**

As described in the main paper (see Discussion section), our population abundance estimates draw on 19 independent surveys, some of which did not follow current best-practice protocols (particularly limited spatial extent of the camera trap array, but also camera positioning and operation).

To assess the sensitivity of abundance and other model estimates to removing some of these surveys, we performed two sensitivity analysis. In the "marginal removal" analysis, we independently dropped each survey and fitted models to capture histories from the remaining 18 surveys. This assesses if there is any single survey that is particularly "influential" on parameter estimates i.e. estimates change substantially if this survey is deleted. In the "cumulative removal" analysis, we first ranked surveys in ascending order of the maximum distance between any pair of cameras used in the survey (this being a proxy for survey size, setting an upper bound on the range of movement that can be detected by recaptures). We then dropped all surveys whose rank was strictly less than $r = 1, \dots 19$, and fitted models to capture histories from the remaining $20 - r$ surveys. For example, $r = 3$ removes the two smallest surveys (which have ranks $r < 3$) and fits models to the remaining 17. The case $r = 1$ uses the full set of 19 surveys.

In all cases two models were fitted: the model preferred by AIC (density a function of elevation, baseline encounter rate a function of camera type) and a null model (spatially uniform density and baseline encounter rate). Models fitted to capture histories with $r > 16$ failed to converge because by this stage the number of detections in the remaining capture histories was extremely small (the three largest surveys made zero, four, and one detections respectively). As removing nearly all surveys was never going to be a feasible option, we did not investigate this further.

Abundance estimates were insensitive to the removal of smaller surveys (Figure A1), changing materially only if surveys spanning less than 50km were excluded. This would constitute a very conservative (approximately $6\sigma$) threshold. For reference, the diameter of a 500km$^2$ home range is 24km. Moreover, increases in abundance estimates if only larger surveys are included did not appear to be a result of smaller surveys underestimating $\sigma$. In fact, estimates of $\sigma$ decreased if smaller surveys were removed (Figure A2). Rather, they appeared to be due to the increasing reliance on two surveys (Khubjerab National Park, 2011; Nagar Hunza Valley, 2018) of relatively high-density areas (note the decrease in abundance estimates if either of these surveys are removed; Figure A1, Marginal removal, points denoted (i) and (ii)). These two surveys are those with the largest number of detected animals (15 in Khubjerab National Park, 9 in Nagar Hunza Valley) and among those with the largest number of detections (24 in Khubjerab National Park, 21 in Nagar Hunza Valley). These areas are unlikely to be representative of the broader region.

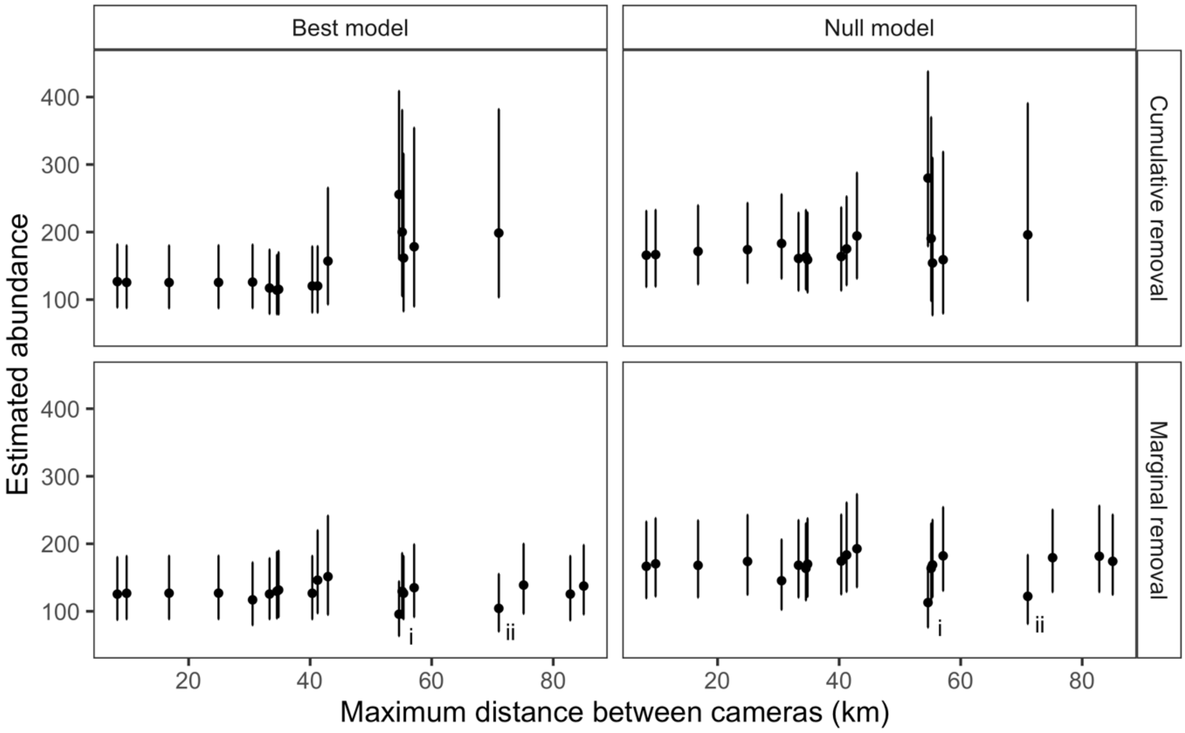


***Figure A1:*** *Estimates of abundance obtained from models fitted to subsets of the 19 surveys used in the main analysis. "Marginal removal" removes each survey independently; "Cumulative removal" removes all surveys with maximum inter-camera distance strictly less than the specified value on the x-axis. Maximum inter-camera distance is used as a measure of the spatial extent of the camera array in each survey. Inset (i) and (ii) denote surveys in Khubjerab National Park and Nagar Hunza Valley respectively (see text for details).*

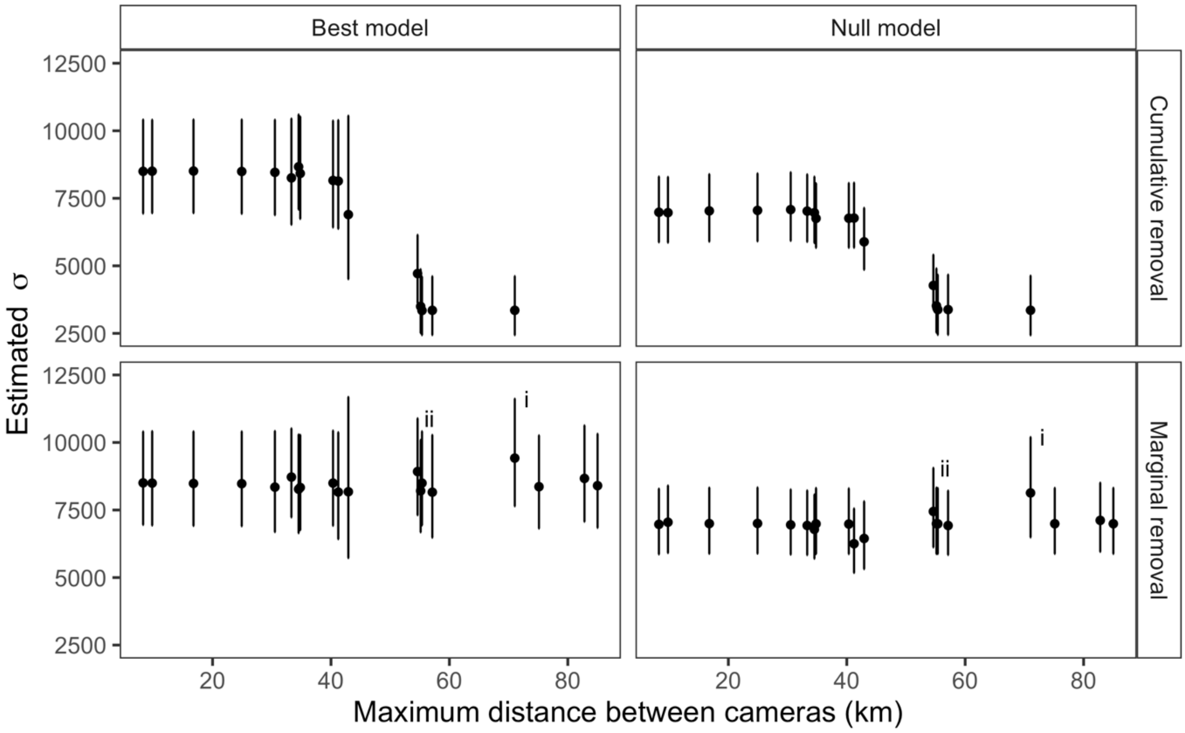


***Figure A2:*** *Estimates of σ obtained from models fitted to subsets of the 19 surveys used in the main analysis. See Figure A1 caption for further details.*
